# Estimated regional white matter hyperintensity burden, resting state functional connectivity, and cognitive functions in older adults

**DOI:** 10.1101/2020.04.14.039065

**Authors:** Abhishek Jaywant, Katharine Dunlop, Lindsay W. Victoria, Lauren Oberlin, Charles J. Lynch, Matteo Respino, Amy Kuceyeski, Matthew Scult, Matthew Hoptman, Conor Liston, Michael W. O’Dell, George S. Alexopoulos, Roy H. Perlis, Faith M. Gunning

## Abstract

**Objective:** White matter hyperintensities (WMH) are linked to deficits in cognitive functioning, including cognitive control and memory; however, the structural and functional mechanisms are largely unknown. We investigated the relationship between estimated regional disruptions to white matter fiber tracts from WMH, resting state functional connectivity (RSFC), and cognitive functions in older adults.

**Design:** Cross-sectional study.

**Setting:** Community.

**Participants:** Fifty-eight cognitively-healthy older adults.

**Measurements:** Tasks of cognitive control and memory, structural MRI, and resting state fMRI. We estimated the disruption to white matter fiber tracts from WMH and its impact on gray matter regions in the cortical and subcortical frontoparietal network, default mode network, and ventral attention network by overlaying each subject’s WMH mask on a normative tractogram dataset. We calculated RSFC between nodes in those same networks.

**Results:** The interaction of estimated regional WMH burden and RSFC in cortico-striatal regions of the default mode network and frontoparietal network was associated with memory retrieval. Models predicting working memory, cognitive inhibition, and set-shifting were not significant.

**Conclusions:** Findings highlight the role of circuit-level alterations at the structural and functional levels in resting state networks that are related to WMH and impact memory retrieval in older adults.

## 1. Objective

Cerebrovascular disease is associated with mood disorders such as late-life depression^1^, and characterized by neurobiological abnormalities like white matter hyperintensities (WMH)^2^. WMH are linked to cognitive decline in older adults; a potential mechanism is their impact on structural connectivity^3^. WMH affect the strength and efficiency of white matter connections, which in turn are associated with slower processing speed^4^ and declines in working memory, episodic memory, and cognitive flexibility^5,6^.

Cognitive control—the ability to maintain task-relevant information in mind, flexibly shift set, and inhibit irrelevant information—is susceptible to cerebrovascular disease and WMH and their impact on structural connectivity. In older adults, cognitive control is linked to structural disconnection in frontal-subcortical circuits arising cerebrovascular disease^7^. These processes have important ramifications for older adults as they may increase risk for late-life mood disorders and treatment non-response^8^ as well as loss of functional independence^9^. Memory dysfunction also occurs in the context of cerebrovascular disease and is associated with WMH-linked disruption in fiber tracts in frontal, temporal, and subcortical regions such as the anterior thalamic tract, the uncinate fasciculus, and the forceps minor^10,11^. Worse memory performance is also associated with depression in older adults^12^.

Cognitive dysfunction in the setting of cerebrovascular disease may also depend on resting state functional connectivity (RSFC). In older adults with small vessel ischemic disease, worse cognitive control is associated with diminished RSFC in the frontoparietal network (FPN), default mode network (DMN), and in subcortical structures^13,14^. White matter lesions are also associated with altered RSFC in the DMN^15^ in mild cognitive impairment. These findings suggest that there are complex and dynamic interrelationships between structural and functional connectivity.

Despite advances in neuroimaging techniques—and much research on the interrelationships among neuroimaging markers of cerebrovascular disease, structural anatomy and RSFC in small vessel disease and mild cognitive impairment—how these interactions contribute to changes in cognitive control and memory in healthy older adults is largely unknown. These interactions may inform the early mechanisms that may underlie cognitive decline in older adults, which can also shed light on circuit-level alterations that may predispose older adults to behavioral and mood disturbances. Understanding the relationship between structure, function, cognition, and vascular disease can also inform targeted and early approaches to promote and maintain cognitive health prior to the onset of late-life psychiatric disorders.

The goal of this study was to investigate how estimated regional WMH burden and RSFC are related to cognitive control and memory in healthy older adults. We used the Network Modification Tool^16^ to estimate the magnitude of disruption to white matter tracts from WMH in the FPN, DMN, and ventral attention network (VAN) and its impact on cognitive control and memory retrieval. We evaluated the relationship between estimated burden from WMH in these networks and RSFC in the same networks. We hypothesized that WMH-associated structural disruption and functional connectivity measures would interact in explaining cognitive performance.

## 2. Methods

### 2.1. Participants

Participants were 58 cognitively-healthy, independent, and community-dwelling older adults aged 60-84 (M=72.9 years, SD=6.02) who were enrolled in a larger trial investigating cognitive control and emotion regulation in late-life depression (ClinicalTrials.gov registration NCT01728194). All participants were English-speaking, non-depressed, and cognitively unimpaired (Mini Mental Status Examination ≥ 26/30). The absence of mild cognitive impairment or dementia was ensured through additional clinical assessment when indicated during the initial screen, including consensus clinical conferences led by board-certified geriatric psychiatrists. Because participants constituted a control group for a larger study, any participant in this sample who reported subjective memory complaints during an initial phone screen was screened out. Participants had no current or past history of major psychiatric illness or neurologic disorder. All participants were recruited through flyers and advertisements. All provided written informed consent, and the study was approved by the Institutional Review Boards of Weill Cornell Medicine and the Nathan Kline Institute. The participants constituted the control group for an analysis on structural connectivity and performance on the Trail Making Test in late-life depression that has been previously published by our group^17^. We extend our previous work by performing analyses of RSFC and relating it to estimated regional WMH burden and to cognitive measures in healthy older adults.

### 2.2. Neuropsychological Assessment

Trained research assistants administered the neuropsychological measures. To assess auditory attention and working memory, we administered the Digit Span subtest (total score of the forward and backward trials) of the Wechsler Adult Intelligence Scale-Fourth edition. The Stroop Color Word Test was used to assess cognitive inhibition and susceptibility to interference from conflicting information. To assess attentional set-shifting we used the Trail Making Test (TMT) and calculated a ratio that isolates the shifting component while accounting for psychomotor processing speed, by dividing the time to completion on TMT-B by the time to completion on TMT-A^18^. We administered the Hopkins Verbal Learning Test-Revised and used the number of words recalled on the delayed recall trial to evaluate memory retrieval.

### 2.3. Neuroimaging

Structural and functional MRI scans were acquired on a 3T Siemens TiM Trio (Erlangen, Germany) scanner that was equipped with a 32-channel head coil at the Center for Biomedical Imaging and Neuromodulation of the Nathan Kline Institute for Psychiatric Research. Structural imaging included high-resolution whole brain images acquired using a 3D T1-weighted MPRAGE and T2-weighted FLAIR images. The acquisition parameters for MPRAGE were: TR=2500ms, TE=3.5ms, slice thickness=1mm, TI=1200ms, 192 axial slices, matrix=256 x 256 (voxel size=1mm isotropic), FOV=256mm, IPAT=2, flip angle=8 degrees. The acquisition parameters for the FLAIR sequence were TR=9000ms, TE=111ms, TI=2500ms, FOV=192 x 256 mm, matrix 192 x 256, slice thickness=2.5 mm, number of slices=64, IPAT=2, flip angle=120 degrees. The final FLAIR resolution (rectangular FOV/matrix) was 1×1 in plane. Functional neuroimaging was a turbo dual echoplanar image (EPI) sequence performed while participants were at rest. Acquisition parameters were repetition time=2500ms, echo time=30ms, flip angle=80 degrees, slice thickness=3mm, 38 axial slices, matrix=72 x 72, 3-mm isotropic, field of view=216mm, integrated parallel acquisition techniques factor=2. Resting-state image acquisition was conducted in a single run that was 6 minutes and 15 seconds in duration, TR=2000ms. Participants were instructed to stay awake with eyes closed. Wakefulness was verified at the end of the scanning sequence by the technician.

Procedures to segment and estimate WMH burden and to estimate the impact of WMH on white matter fiber tracts have been previously described in detail^17^. Figure 1 illustrates the flow of neuroimaging analysis and statistical procedures. In brief, two raters performed a visual rating using operational criteria of the Age-Related White Matter Change scale^19^. Inter-rater reliability was strong (Intraclass Correlation Coefficient=.95). FSL^20^ and the BIANCA program were subsequently used to segment WMH lesions and create WMH masks. WMH lesions smaller than 3 voxels were removed. A visual check was performed on each individual WMH mask and minimal manual adjustments were made. Each final WMH mask was nonlinearly registered to MNI space and binarized.

**Figure 1.**
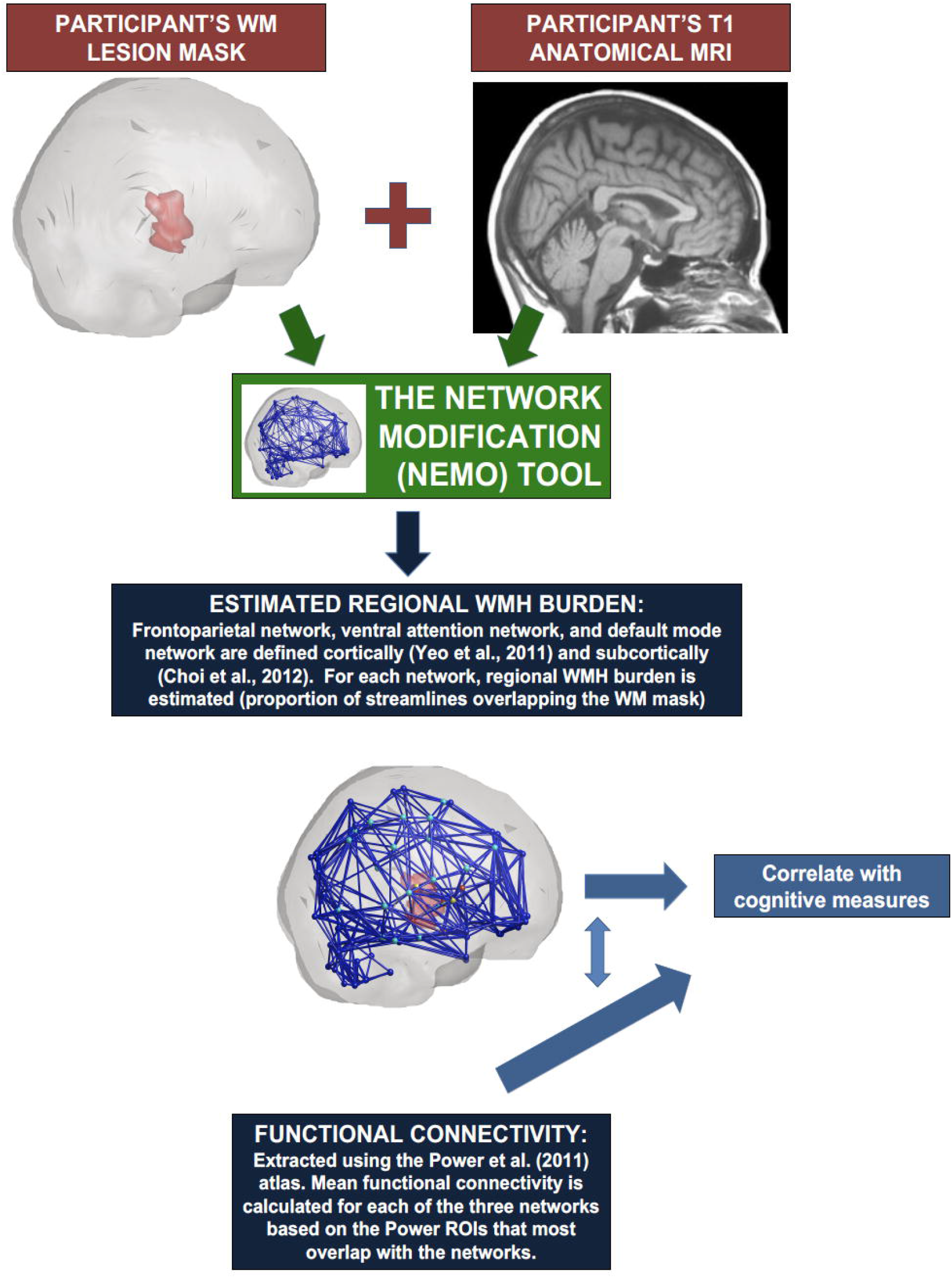
Flowchart of neuroimaging analysis procedures and statistical procedures for the primary analyses.

Regional WMH burden was estimated using the Network Modification Tool, which has previously been used in aging populations^21^. Each WMH mask was overlaid on a normative sample of 420 healthy participants’ tractograms from the Human Connectome Project. The program estimates the disruption to white matter fiber tracts and the effects on gray matter regions, incorporating the potential distal effects of white matter lesions on gray matter, by removing those tracts passing through a subject’s WMH mask. This procedure was conducted using a combined atlas of the Yeo et al^22^ 7-network cortical parcellation and the Choi et al^23^ 7-network corresponding striatal parcellation. We were primarily interested in the FPN, DMN, and VAN, the latter because it includes nodes of the salience network such as the dorsal anterior cingulate that are implicated in cognitive control. For our analysis, we evaluated the proportional loss (due to WMH mask overlap) in tracts cortically in the three networks, and between the cortex and striatum because of the known association between cortico-striatal connectivity and cognition. The proportional loss of tracts due to the WMH mask was calculated for each network (cortical and cortico-striatal separately) by dividing the number of streamlines in that subject’s modified connectome by the mean number of streamlines in that region pair from the 420 healthy control tractogram set. Values range from 0 to 1, with higher values indicating greater estimated regional WMH burden.

To extract corresponding RSFC values, each subject’s EPI images were motion corrected using FSL’s MCFLIRT, linearly registered to the anatomical T1 scan, and normalized to the MNI 152 template. Additional preprocessing including slice-timing correction and spatial smoothing (6mm FWHM) was performed in AFNI^24^. Further removal of motion artifacts was performed using ICA-AROMA^25^. Motion was regressed out (demeaned and first derivative), artifacts were further removed using AFNI’s *anaticor*, and the time-series was bandpass filtered. Time-series data were extracted using a previously published parcellation of 277 functional nodes^26^ that includes the 264 nodes from the Power atlas^27^ combined with 13 additional nodes in the caudate, amygdala, hippocampus, nucleus accumbens, subgenual anterior cingulate, locus coeruleus, ventral tegmental area, and the raphe nucleus. Each subject’s 277×277 matrix was Fisher r-to-Z transformed. We subsequently extracted the modified Power atlas ROIs overlapping with the combined Yeo-Choi parcellation by calculating the Dice similarity coefficient for each network (FPN, VAN, DMN cortical and striatal). The 10 functional nodes in each network that had the largest Dice coefficient > 0 (i.e. had > 1 voxel overlapping the structural ROI) were retained. We averaged the RSFC for the nodes within each network to extract a per-network RSFC score. Note that we elected not to use the same 7-network parcellation for RSFC because current recommendations suggest using functionally-based parcellations with much more fine-grained nodes for fMRI data^28^.

### 2.4. Statistical Analysis

Statistical analyses were conducted using IBM SPSS Statistics version 25 (IBM, Armonk, NY), Matlab, and RStudio. Analyses focused on the FPN, DMN, and VAN from the combined Yeo-Choi parcellation as described above. We computed Spearman rank-order correlations (because of the possibility of a non-linear relationship) to evaluate the relationship between estimated regional WMH burden and RSFC in each network, cortical and cortico-striatal separately. We used a conservative alpha of .01 given 36 correlations in the matrix, rather than a Bonferroni correction, to best balance the risk of type I and type II error and to ensure that we could detect correlations of modest strength. To further evaluate the robustness of these relationships, we compared model fit using the likelihood ratio test in predicting RSFC from regional WMH burden beyond the effect of age, education, and gender (see Supplemental Digital Content 1).

We conducted a multivariate general linear model with alpha=.05 to determine how estimated regional WMH burden and functional connectivity were associated with cognitive performance. Outcome variables were Digit Span, Stroop Interference, TMT B/A ratio, and HVLT-R Delayed Recall. As predictors, we entered as main effects: (1) estimated regional WMH burden within the cortical FPN, DMN, and VAN; (2) cortico-striatal estimated regional WMH burden within the FPN, DMN, and VAN; (3) cortico-cortical RSFC within the FPN, DMN, and VAN; and (4) cortico-striatal RSFC within the FPN, DMN, and VAN. We entered as interaction effects— which was the primary test of our hypothesis of structure-function interactions—the estimated regional WMH burden x RSFC for each network (i.e., cortico-cortical estimated regional WMH burden x RSFC in the FPN, cortico-striatal estimated regional WMH burden x RSFC in the FPN, cortico-cortical estimated regional WMH burden x RSFC in the DMN, cortico-striatal estimated regional WMH burden x RSFC in the DMN, cortico-cortical estimated regional WMH burden x RSFC in the VAN, cortico-striatal estimated regional WMH burden x RSFC in the VAN). All predictors were Z-normalized. Age, education, and gender were entered as covariates.

## 3. Results

### 3.1. Demographic and clinical variables

Table 1 displays the mean, SD, and ranges for age, education, and performance on the dementia screening assessments (Mini Mental Status Examination) and neuropsychological outcome measures for our sample. The ratio of male:female participants is also provided. Figure 2 shows the WMH distribution in the sample.

**Figure 2:**
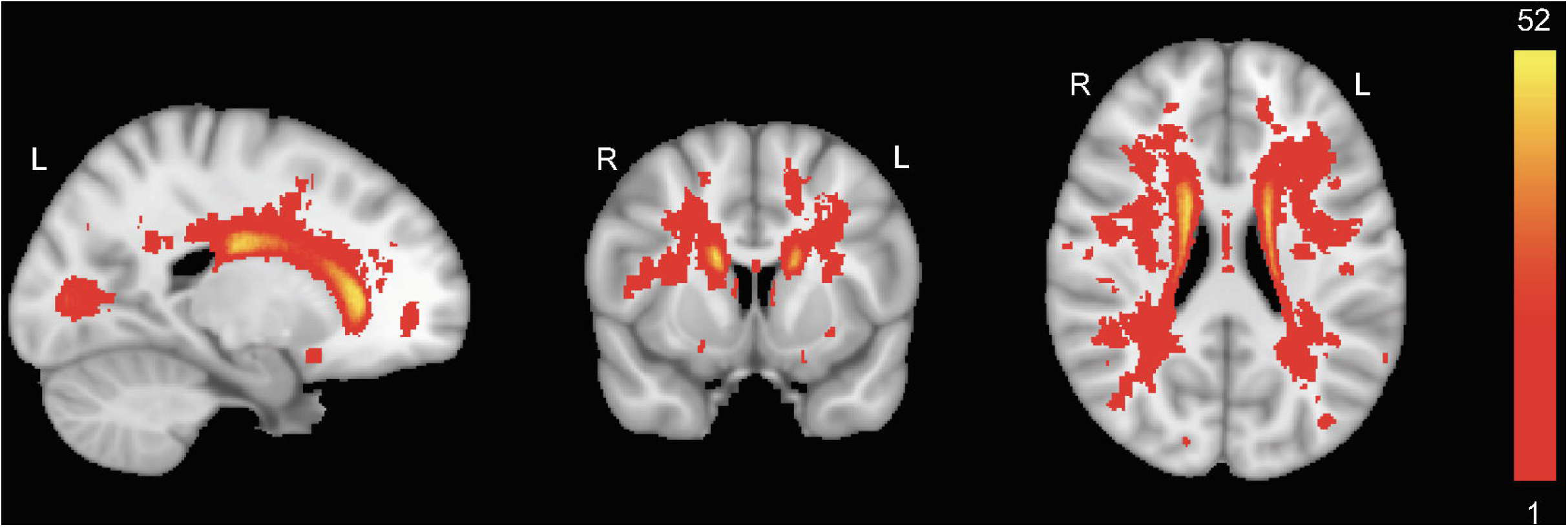
Heatmap depicting the distribution of white matter hyperintensities (WMH) in the N=52 participants analyzed in the final sample. The heatmap was created by concatenating all participants’ binary lesion masks and thus values range from 0 to 52.

**Table 1.**
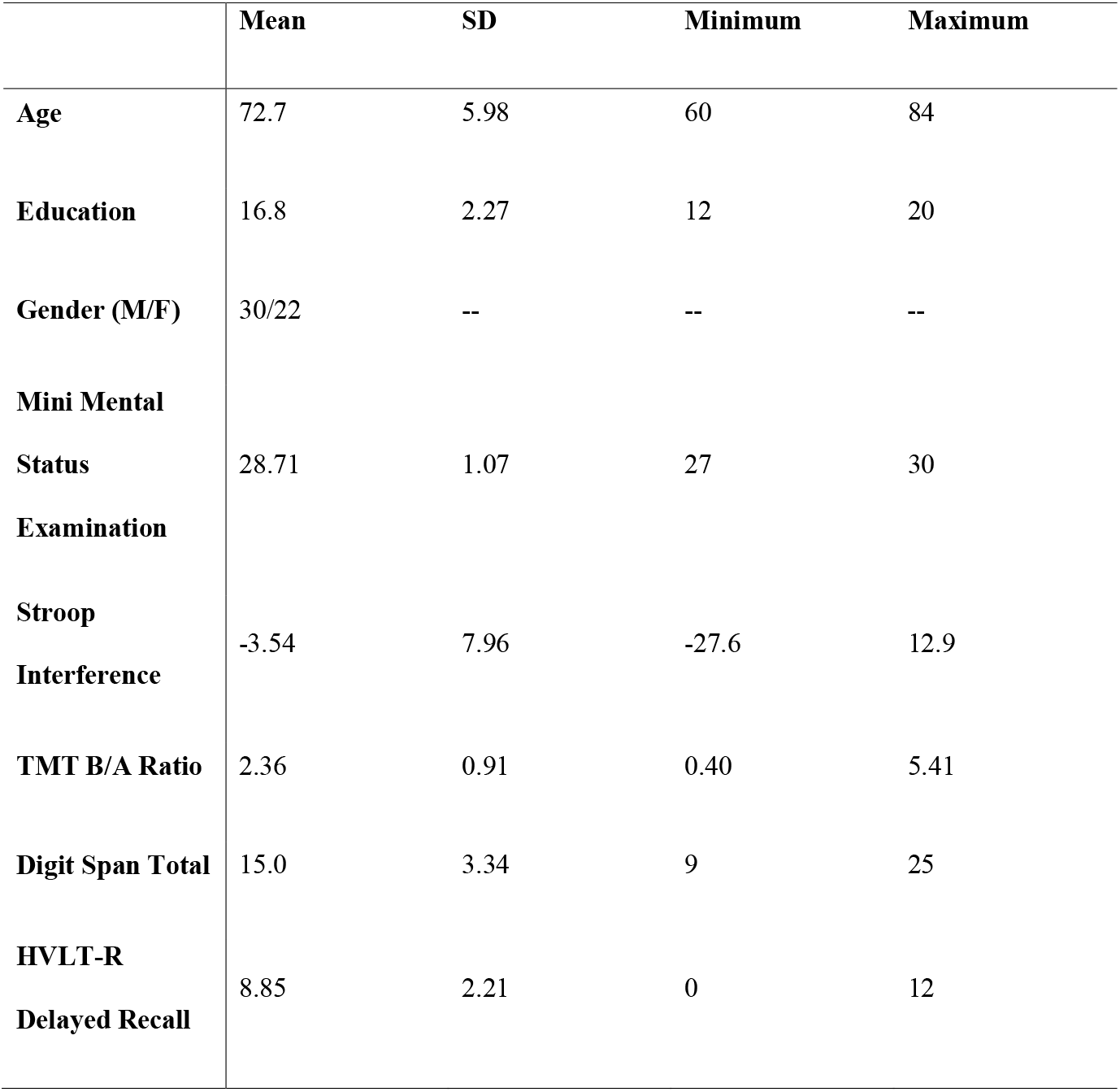
Demographic and clinical characteristics of the analyzed study sample (N=52).

### 3.2. Correlations between estimated regional WMH burden and RSFC

Table 2 shows the correlations between estimated regional WMH burden and RSFC for the FPN, VAN, and DMN, both cortically and between the cortex and striatum. Six participants were excluded from the correlation analyses and the multivariate general linear model (below) due to missing data, resulting in a final analyzed sample of N=52. There were significant negative correlations (p<.01) between RSFC in the cortical DMN and estimated regional WMH burden in the cortico-cortical FPN and cortico-striatal FPN. There were significant negative correlations between RSFC in the cortico-cortical VAN and estimated regional WMH burden in the cortico-cortical VAN. Of these correlations, only the relationship between estimated regional WMH burden in the cortico-striatal FPN and RSFC in the cortico-cortical DMN improved model fit beyond using age, education, and gender to predict RSFC (Supplemental Digital Content 1).

**Table 2.** Spearman rank-order correlations between estimated regional WMH burden (proportion loss of connections) and resting state functional connectivity (RSFC; Fisher r to z transformed) in networks of interest.

| **Estimated Regional WMH Burden:** | **RSFC Cortical VAN** | **RSFC Cortical FPN** | **RSFC Cortical DMN** | **RSFC Cortico-Striatal VAN** | **RSFC Cortico-Striatal FPN** | **RSFC Cortico-Striatal DMN** |
| --- | --- | --- | --- | --- | --- | --- |
| **Cortical VAN** | -.36, p=.009 | -.24, p=.09 | -.29, p=.04 | .09, p=.51 | -.23, p=.10 | -.18, p=.21 |
| **Cortical FPN** | -.25, p=.08 | -.15, p=.30 | -.38, p=.005 | -.11, p=.43 | -.19, p=.18 | -.16, p=.27 |
| **Cortical DMN** | -.30, p=.03 | -.15, p=.30 | -.35, p=.01 | -.13, p=.35 | -.18, p=.20 | -.09, p=.55 |
| **Cortico-Striatal VAN** | -.34, p=.01 | -.31, p=.03 | -.33, p=.02 | -.13, p=.35 | -.27, p=.05 | -.14, p=.33 |
| **Cortico-Striatal FPN** | -.29, p=.04 | -.11, p=.45 | -.36, p=.008* | -.14, p=.32 | -.20, p=.16 | -.06, p=.66 |
| **Cortico-Striatal DMN** | -.33, p=.02 | -.08, p=.59 | -.27, p=.06 | -.04, p=.76 | -.13, p=.36 | -.13, p=.36 |

Note. VAN = Ventral Attention Network; FPN = Frontoparietal Network; DMN = default mode network. To reduce the probability of a Type I error, we interpret as significant correlations with $p < .01$ . \*Denotes relationship in which estimated regional WMH burden improved model fit in predicting RSFC beyond age, education, and gender (see Supplement 1).

### 3.3. Associations between estimated regional WMH burden, RSFC, and cognition

Results of the omnibus test of significance for the multivariate general linear model indicated a significant effect on HVLT-R Delayed Recall, *F*(24)=3.43, *p*=.001, adjusted *R^2^*=.50. The omnibus test was not significant for Stroop Interference, *F*(24)=.90,*p*=.59, adjusted *R^2^*=-.04; Digit Span, *F*(24)=1.37, *p*=.21, adjusted *R^2^*=.13; or TMT B/A, *F*(24)=.22, *p*=1.0, adjusted *R^2^*=-.47). We did not evaluate the significance of individual predictors for these non-significant outcomes.

For the outcome variable of HVLT-R Delayed Recall, our primary analysis was the interaction between regional WMH burden and RSFC. There were significant interaction effects of estimated regional WMH burden x RSFC in the cortico-striatal FPN (β=-.89, SE=.31, *t*=-2.89, *p*=.007), cortico-striatal DMN (β=.68, SE=.31, *t*=2.18, *p*=.04), and cortico-cortical DMN (β=1.67, SE=.79, *t*=2.11, *p*=.04). For the interaction between estimated regional WMH burden and RSFC in the cortical DMN, there was a negative relationship between RSFC and memory retrieval, although the slope of that relationship was weaker in individuals with low estimated regional WMH burden (Figure 3a; Figure 4). For the interaction between estimated regional WMH burden and RSFC in the cortico-striatal DMN, higher RSFC in the DMN was associated with better memory retrieval for individuals with high estimated regional WMH burden; lower RSFC was associated with better memory retrieval for those with low estimated regional WMH burden; and there was minimal relationship between RSFC and memory retrieval for those with moderate estimated regional WMH burden (Figure 3b; Figure 4). For those with high estimated regional WMH burden in the cortico-striatal FPN, lower RSFC was associated with better memory retrieval, while the opposite was true of those with low and moderate estimated regional WMH burden (Figure 3c; Figure 4). Though not of primary interest in evaluating our hypotheses, significant main effects included education (β=.30, SE=.12, *t*=2.45, *p*=.02), RSFC in the cortical FPN (β=-2.68, SE=1.06, *t*=-2.53, *p* = .02), and RSFC in the cortico-striatal FPN (β=.76, SE=.34, *t*=2.20, *p*=.04).

**Figure 3.**
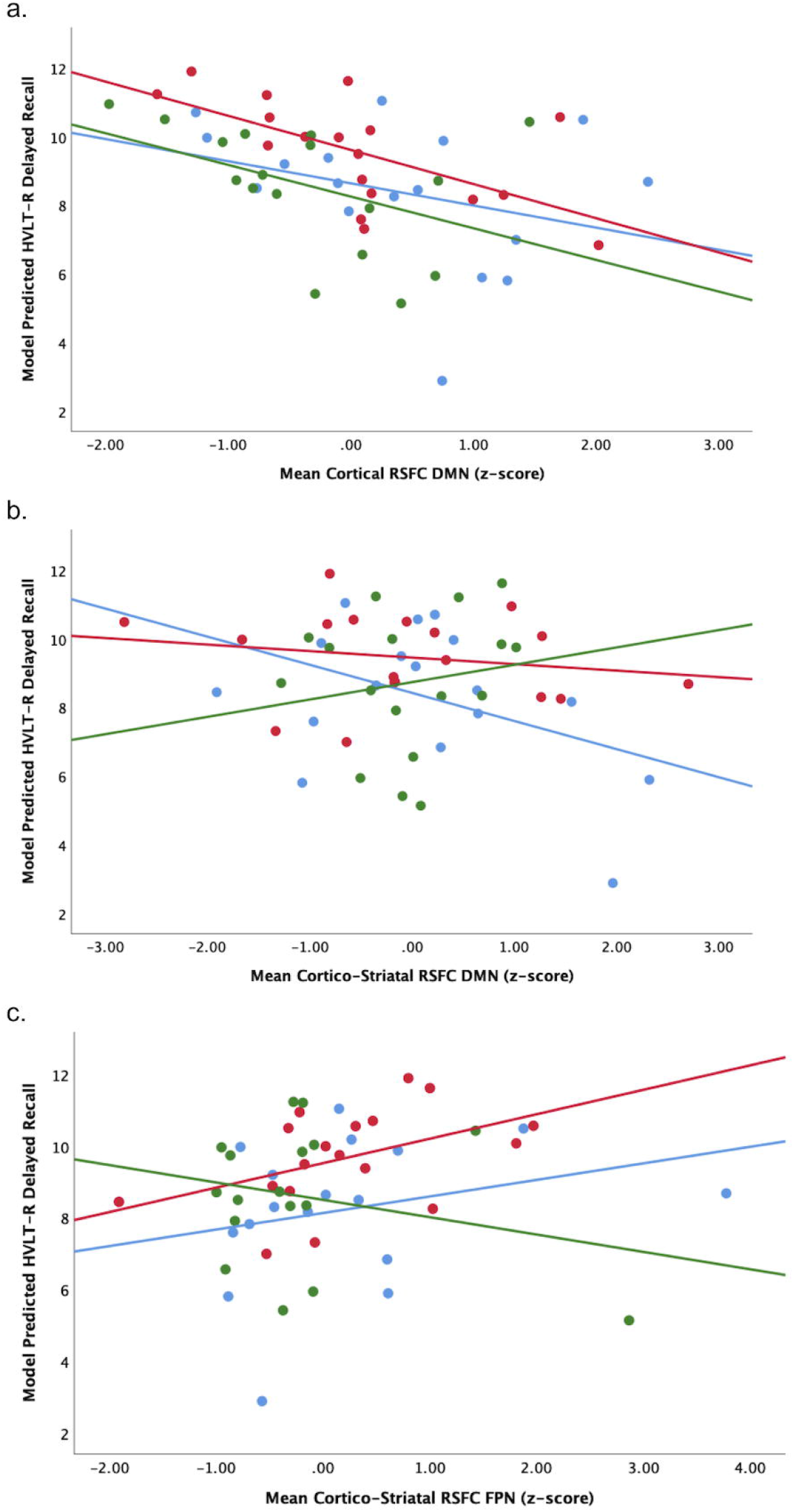
Scatterplots (with best fit lines) depicting the relationship between HVLT-R Delayed Recall performance predicted by the general linear model, estimated regional white matter hyperintensity burden, and resting state functional connectivity (RSFC) in the (a) cortical default mode network (DMN); (b) cortico-striatal DMN; and (c) cortico-striatal frontoparietal network (FPN). For visualization, estimated regional white matter hyperintensity burden was divided into three equal groups: “low” (blue dots), “moderate” (red dots), and “high” (green dots) burden. RSFC values are normalized to z-scores as in the general linear model.

**Figure 4.**
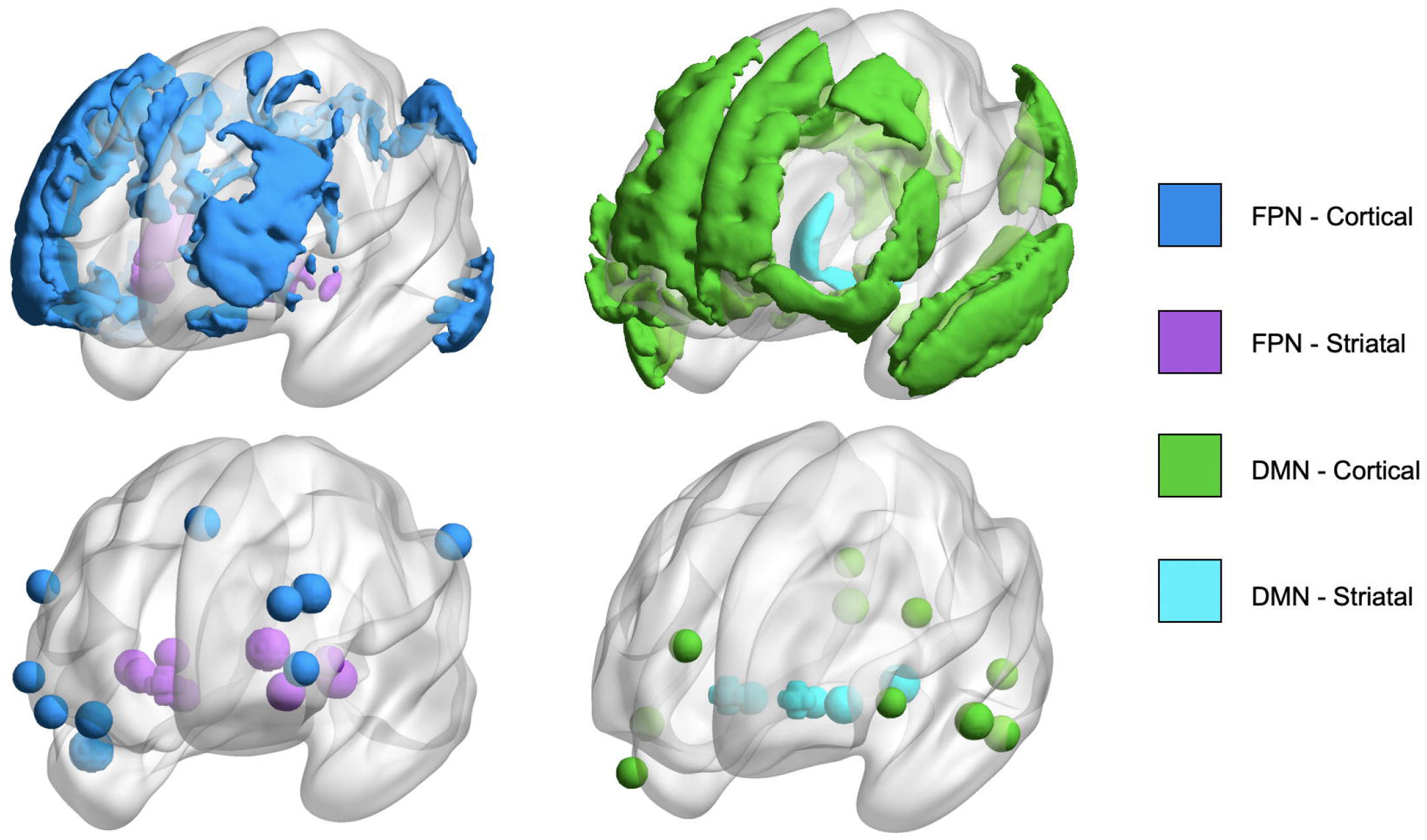
Glassbrains depicting the frontoparietal network (FPN; top left) and default mode network (DMN; top right) as defined by the combined Yeo-Choi atlas and used to calculate the estimated regional white matter hyperintensity burden. Glassbrains also depict functional nodes in the FPN (bottom left) and DMN (bottom right) as defined by the Power atlas and used to calculate resting state functional connectivity.

## 4. Discussion

We found that in cognitively-healthy older adults, (1) there are modest associations between estimated regional WMH burden and RSFC within and across networks, with WMH burden in the FPN and RSFC in the DMN having a relatively robust association; and (2) the relationship between estimated regional WMH burden and resting state functional connectivity predicts memory retrieval, but not cognitive inhibition, attention/working memory, or set-shifting. Specifically, estimated regional WMH burden interacts with RSFC in cortico-cortical and cortico-striatal regions of the FPN and DMN, to predict memory retrieval. These findings extend prior work relating white matter lesions and RSFC in small vessel disease and mild cognitive impairment. Our findings indicate that alterations in structural and functional circuitry from WMH are both associated with select aspects of cognitive functioning in healthy older adults.

We found that structure-function relationships may differ by network in cognitively-healthy older adults without a diagnosed cerebrovascular disorder. These findings suggest that greater estimated regional WMH burden is associated with an age-related decline in RSFC within and between networks, though the magnitude of the correlation coefficients were small-moderate. We found a relatively robust negative association between cortico-striatal estimated regional WMH burden in the FPN and RSFC in the cortico-cortical DMN. This finding is consistent with the known dense connections that exist between networks^29^. It is also consistent with observations that RSFC decreases in functional networks, including the DMN, in normal aging^30^, small vessel ischemic disease^13^, and stroke^31^. Although RSFC in the FPN was not associated with structural disruption from WMH, there is suggestion that structure-function coupling within this network may weaken in older adults^32^.

Our examination of estimated regional WMH burden replicates prior studies demonstrating a relationship between disruptions in white matter pathways and cognitive decline in older adults^4,6^, and the effects of WMH on distal white matter tracts and cognition^33^, and extends them to identify interactions with RSFC. We found that the relationship between estimated regional WMH burden and RSFC in the FPN and DMN predicted memory retrieval. This finding is consistent with the known role of the DMN in episodic memory^34^ and the importance of the role of executive control processes subserved by the FPN in facilitating effortful search and retrieval from long-term memory^35^.

The interactions observed in the DMN and FPN and memory retrieval indicates that in individuals with high estimated regional WMH burden in the cortico-striatal DMN, RSFC was positively associated with memory retrieval, while RSFC was negatively associated with memory retrieval in those with low estimated regional WMH burden. This pattern suggests that functional connectivity in the DMN may support behavioral performance in a “compensatory” manner in the face of increasing WMH-related structural disruption. Increases in functional connectivity may be protective against the negative effects of white matter lesions^36^. Our result is also similar to the finding that greater functional connectivity between the medial prefrontal cortex and inferior parietal cortex is associated with better episodic memory in older adults with greater gray matter atrophy^37^.

Consistent with the known role of functional activation in the FPN during memory retrieval processes, increasing RSFC in the cortico-striatal FPN was positively associated with better memory retrieval in those with low and moderate levels of estimated regional WMH burden. However, at high levels of WMH burden, increasing RSFC in the FPN was associated with worse memory performance. This pattern may represent an unsuccessful or maladaptive compensatory response, reflected in an overreliance on the FPN. A similar dissociation between functional activity in the DMN and FPN at high levels of WMH-related structural disruption has been observed recently in multiple sclerosis in which cognitive deficits were linked to decreased DMN activity and increased FPN activity^38^. That cortico-striatal connectivity in the FPN and DMN was implicated may be because the WMH in our sample were frequently in periventricular regions.

Contrary to our initial hypothesis, we did not find significant associations with cognitive control. The lack of association with cognitive control may be related to the assessment measures themselves, as the neuropsychological measures we used may not have been sensitive or comprehensive enough to detect subtle age-related decline in cognitive control functions. In addition, subjective cognitive decline may be a marker of subtle deficits prior to objectively observable cognitive impairment, thus, it is possible that estimated regional WMH burden and RSFC may have predicted subjective complaints in cognitive control functions.

Limitations include the length of the resting state scan (6 minutes and 15 seconds), which is of short duration compared to current standards. Given that reliability tends to increase with longer durations replication of our RSFC findings with longer scan times is warranted. Our age range was relatively restricted and future studies would benefit from including a broader range of ages and comparing our findings to middle-aged and old-old adults. The normative tractograms used by the Network Modification Tool are drawn from relatively young adults in the Human Connectome Project and the discrepancy in age with our sample may have induced additional error. However, this weakness may be partially mitigated by studies demonstrating that structural disconnection maps were similar in younger adults versus older adults^39^ and a strong correlation (r=.87) between disconnection maps that use age-matched reference tractograms and young adult tractograms from the Human Connectome Project^40^. Our measure of estimating regional burden from WMH takes an “all or none” approach (presence or absence of a streamline) which likely oversimplifies the process of white matter injury and reorganization; nonetheless, this approach can be useful in clinical populations in determining inferred disconnection in a cost-efficient manner when diffusion MRI is not feasible.

### 4.1. Conclusion

We demonstrate that in cognitively-healthy older adults, estimated regional WMH burden is associated with a decline in resting state functional connectivity in the DMN, and that the interaction between resting state functional connectivity and estimated regional WMH burden in cortical and subcortical regions of the frontoparietal and default mode networks are both associated with memory retrieval. Our findings highlight the role that alterations in the structural and functional connectome both play in older adults in the setting of cerebrovascular processes. Early, targeted approaches directed towards the FPN and DMN may promote and maintain cognitive health in older adults, prior to the onset of mild cognitive impairment or late-life mood disorders.

## Acknowledgements

We thank Cristina Pollari for assistance with database management, Naib Chowdhury for help acquiring behavioral data, Keith Jamison for assistance in implementing the Network Modification Tool, and Elvisha Dhamala for help in creating the figures. We are grateful to the staff of the Center for Biomedical Neuroimaging and Brain Modulation at the Nathan Kline Institute for Psychiatric Research for assistance with neuroimaging data collection. This work was supported by the National Institute of Mental Health (NIMH) grants R01 MH097735 (Gunning) and T32 MH019132 (Alexopoulos). The sponsor did not have any role in the study design, the acquisition, analysis, or interpretation of the data, the writing of the report, or the decision to submit the manuscript for publication.

## Author Contributions

AJ: conceptualization, methodology, formal analysis, data curation, writing, and visualization. KD: conceptualization, methodology, analysis validation, and writing. LWV: conceptualization, methodology, data curation, writing, and visualization. LO: conceptualization, writing; CJL: conceptualization, writing; MR: methodology, conceptualization, software implementation; data curation, formal analysis; AK: methodology, software development, conceptualization, writing; MS: writing; MJH: writing; CL: supervision; MWO: writing; GSA: writing, funding acquisition; RHP: formal analysis, writing; FMG: conceptualization, methodology, writing, supervision, funding acquisition.

## List of Supplemental Digital Content

Supplemental_Digital_Content_1.docx

